# Electroantennography Responses of African Meliponine Bee Species (Hymenoptera: meliponini) to Hetero-Specific Trail Pheromones: Glandular Sequestration of Terpenyl Esters

**DOI:** 10.1101/2024.03.19.585691

**Authors:** O. Bobadoye Bridget, T. Nganso Beatrice, N Kiatoko

**Affiliations:** International Center of Insect Physiology and Ecology, P.O. Box 30772-00100 Nairobi, Kenya; Forestry Research Institute of Nigeria, P.M.B 5047, Ibadan, Oyo state, Nigeria

**Keywords:** Antennal response, exocrine glands, trail pheromones, odor

## Abstract

Preliminary studies of electroantennographic responses of African meliponine bees to trail pheromones were performed using four meliponine species; *Hypotrigona gribodoi, Meliponula ferruginea* (black), *Meliponula ferruginea* (reddish brown) and *Melipona bocandei*. Experiments with *Meliponula ferruginea* (black) and (reddish brown) revealing antennae responses to pentane extracts containing the nasonov gland and tarsal glands respectively. Because the glandular origin of pheromone marks deposited by African meliponine bee’s species has not yet been investigated, we first confirmed if these species carry out scent marking and recruitment behavior at floral sources. Secondly we tested if either nasonov or tarsal gland secretions elicited trail-following behavior in newly recruited bees by means of chemical and electrophysiological analyses as well as with bio-assays testing both natural extracts and synthetic pheromone compounds from both glands. A significantly higher proportion of foragers from the four species were attracted and recruited additional foragers to floral resources baited with natural extracts from their own tarsal and nasonov glands. The dominant compound from postpharyngeal gland extracts detected by the chemo-receptors on the foragers’ antennae from these four species belongs to the chemical class of terpenoid esters, with (*E)*-β Farnesene constituting a dominant part of this trail pheromone in all four species. This terpenoid could potentially be a singular olfactory cue with dual functionality for resource partitioning and competition avoidance between con-specific and hetero-specific foragers on floral resources, which commonly occurs when foragers from different colonies are scouting for food sources in overlapping foraging areas.

## INTRODUCTION

Meliponine bees of the hymenopteran group *Meliponini* exhibit complex communication and behavioral patterns, and much of that communication is facilitated using specialized gland secretions. The study of pheromones in African meliponine bees is still very much at its infancy stage, however, the chemical structures of trail pheromonal compounds have only been elucidated for a small number of meliponine species of neo-tropical origin, but none of African origin till date. The most important exocrine glands in meliponine bees found to be implicated in pheromone production come from body regions which include the head region where the postpharyngeal (PPG) and mandibular glands (MG) are found; the thorax which houses the thoracic salivary glands (TG); and the abdomen where the dufour gland (DG), nasonov (NG) and epidermal glands (EG) are found.

Mandibular gland secretion of meliponine bees has been the most extensively studied gland in honey bees, because it appears to play an important role in chemical communication (Kerr *et al*., 1963). Cephalic secretions of a large number of Apidae species have also been studied extensively as it plays an important role in the behavioral communication of the honey bee (Engels *et al*., 1993, 1997; Francke *et al*., 2000). Understanding the chemical components of other glands will help unravel behavioral patterns which these pheromones support. This has led to some early studies with pure synthetic chemicals which were conducted to understand their influence on the behavior of meliponine bees (Weaver *et al*., 1975; Smith & Roubik, 1983).

Honeybees are known to detect odors using olfactory receptor cells located in their antennae (Lacher, 1964; Vareschi, 1971; Williams *et al*., 1982), and this has been experimentally confirmed using the GC-EAD. The electroantennography detection technique (GC-EAD) which measures the summation of receptor potentials from responding cells in insect antennae (Kaissling, 1971) has supported most studies on chemosensory research especially in insects. It has been used to identify active pheromone components in a mixture of volatile compounds extracted from insects, and has played a vital role in the identification of most pheromones from the insect order.

The recognition of active compounds in GC-EAD studies has proven to help in identifying active compounds capable of eliciting a positive response recognized by antennal receptors which are implicated in pheromonal communication. Meliponine foragers are speculated to use a variety of communication mechanisms to effectively recruit workers of a colony to collect sufficient amounts of food to nourish the entire nest population. Mechanisms used to convey such information include thoracic vibrations and trophallaxis within the nest; footprint secretions and pheromone marks deposited in the field, or a combination of these signals and cues.

Out of all these behavioral mechanisms, there has been numerous discrepancies about the origin of trail pheromone production from the head, thorax, abdomen and leg regions of meliponine bees. This study was carried out to test the hypothesis that a) African meliponine bees carry out scent marking behavior at floral sources and can effectively recruit other foragers to the selected food source b) pheromones responsible for scent marking behavior originate from the nasonov gland but maybe deposited by the tarsal glands upon transfer. Because the glandular origin of pheromone marks deposited by African meliponine bee’s species has not yet been investigated, we first confirmed if these species carry out scent marking and recruitment behavior at food sources. Secondly we tested if either nasonov or tarsal gland secretions elicited trail-following behavior in newly recruited bees by means of chemical and electrophysiological analyses as well as with bio-assay observations evaluating both natural extracts and pure synthetic pheromone compounds.

## METHODS AND MATERIALS

Behavioral experiments were conducted between April and September, 2017 at the laboratory of the behavioral and chemical ecology unit of the International Centre of Insect Physiology and Ecology (*icipe*), Duduville campus (1º 17’S, 36º 49’E) in Nairobi, Kenya. In February 2017, 12 colonies which had been sourced from Taita taveta county (03° 20’ S, 38° 15’ E) were transported to the meliponary section of the International center for insect physiology and ecology (*icipe*) where they were further stabilized and maintained throughout the experimental period. Three colonies each of *Plebeina hildebrandti, Meliponula ferruginea (black), Hypotrigona gribodoi and Hypotrigona ruspolii* used in the experiments were queen right and estimated to be approximately similar in size and fitness, having similar numbers of workers (> 500) individuals. They were placed at a distance of 1m from each other and left to forage freely on nearby vegetation throughout the experimental period.

### Behavioral experiment 1: Scent marking behavior on food resources

On the day when each bio-assay was to be conducted, a total of 12 marked artificial feeders were randomly baited with different artificial food sources (nectar, pollen and water) respectively. The experimental setup and procedure followed the method for scent trail bio-assays described in Jarau *et al*., (2006). Foragers from each colony were gradually trained over a period of two months (February-March) to collect unscented resources from individually marked artificial feeders prior to conducting these observational bio-assays. Observations were made between 09:30 and 15:00, for twenty-five minutes per hour on each feeder. Throughout all observations, the species identity, number of bees landing on each baited feeder and time of collection was recorded. Most importantly the observation of scent marking behavior was observed and confirmed when bees raised their abdomens at an angular length in the air while simultaneously fanning their wings or rubbed their abdomen against their tarsal region (metatarsus/tarsus) (Fig 2) after landing on the feeders.

### Extraction of glands for Bio-assays

Both nasonov and tarsal glands from five foragers of each species returning from foraging bouts were collected from each colony. Bees were collected and immobilized by placing on ice for ∼20 minutes. Prior to gland extraction, hind legs bearing any substance (pollen, nectar or resin) which could be a source of contamination were excised. Gland extraction procedure and concentration of gland extracts was routinely carried out as described by Jarau *et al*., (2006). Glands were dissected in saline solution under a stereo microscope by carefully separating them from any tissue other than the targeted glandular epithelia and thereafter soaked in pentane for 24 hours at room temperature (24°C) (Fig 1). For all extracts, the amount of pentane was adjusted to 100μl per pair of glands (e.g., 10 nasonov/tarsal glands in 500 μL pentane). This is to ensure that 100μl of the pooled extracts corresponded to the gland content of one individual bee (one bee equivalent). 12 extracts (nasonov/tarsal glands) were prepared from each of the four species along with a control (99% pure pentane) in the same manner. These extracts were stored in -20°C until ready to use for behavioral bio-assays.

**Figure 1:**
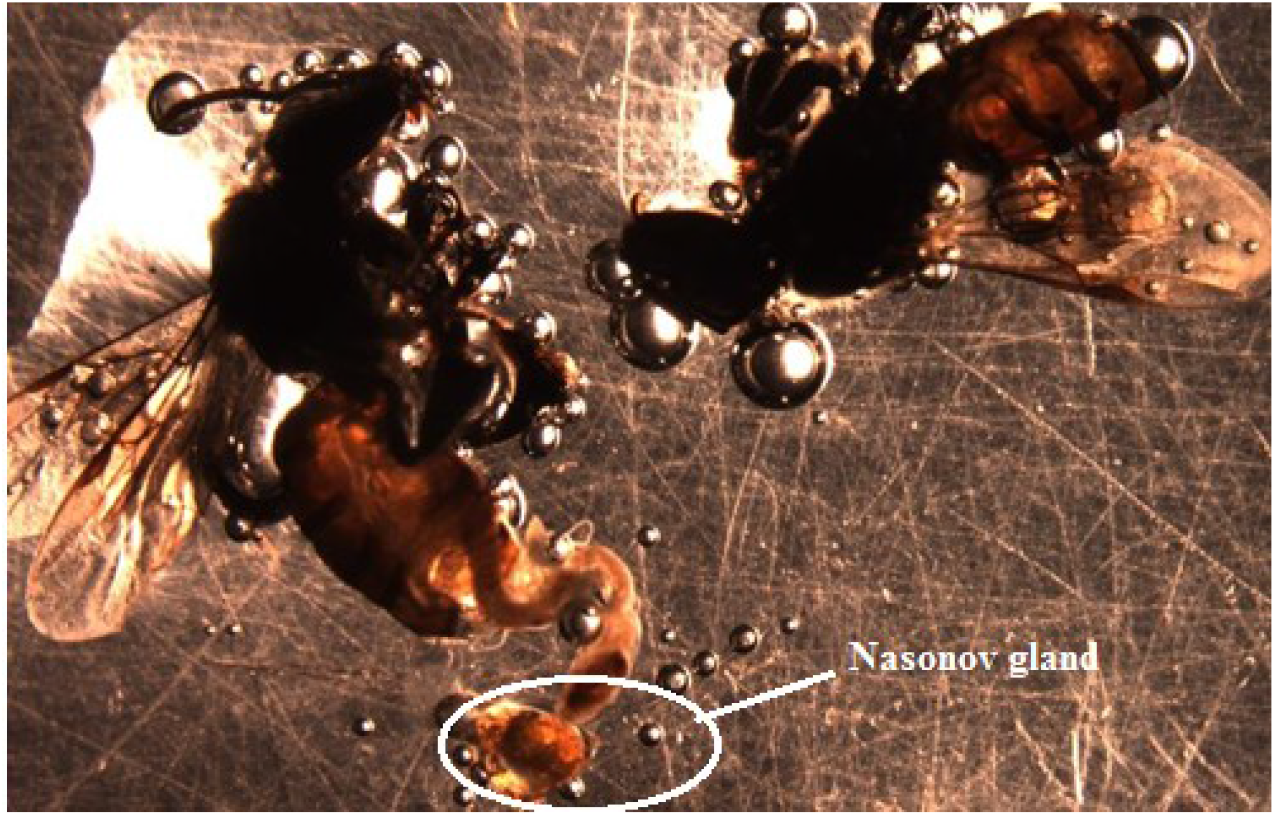
Excised abdominal region containing the nasonov gland (glandular epithelia) from *H. ruspolii* prior to solvent extraction.

### Behavioral experiment 2: Scent marking behavior on food resources baited with natural gland extract

Experiments were carried out on food sources (nectar, pollen and water) baited with either nasonov or tarsal glands respectively from the four species. Approximately 10μl of gland extract were applied on the landing base of each feeder. Observations were made between 09:30 and 15:00, for 25 minutes per hour on each feeder, for 30 days. Throughout all observations, the species identity, number of bees landing on each baited feeder and time of collection was recorded.

Most importantly the observation of scent marking behavior was observed and confirmed to be initiated when bees raised their abdomens at an angular length in the air while simultaneously fanning their wings or rubbed their abdomen against their tarsal region (metatarsus/tarsus) (**Fig 2**) after landing on the feeders.

**Figure 2:**
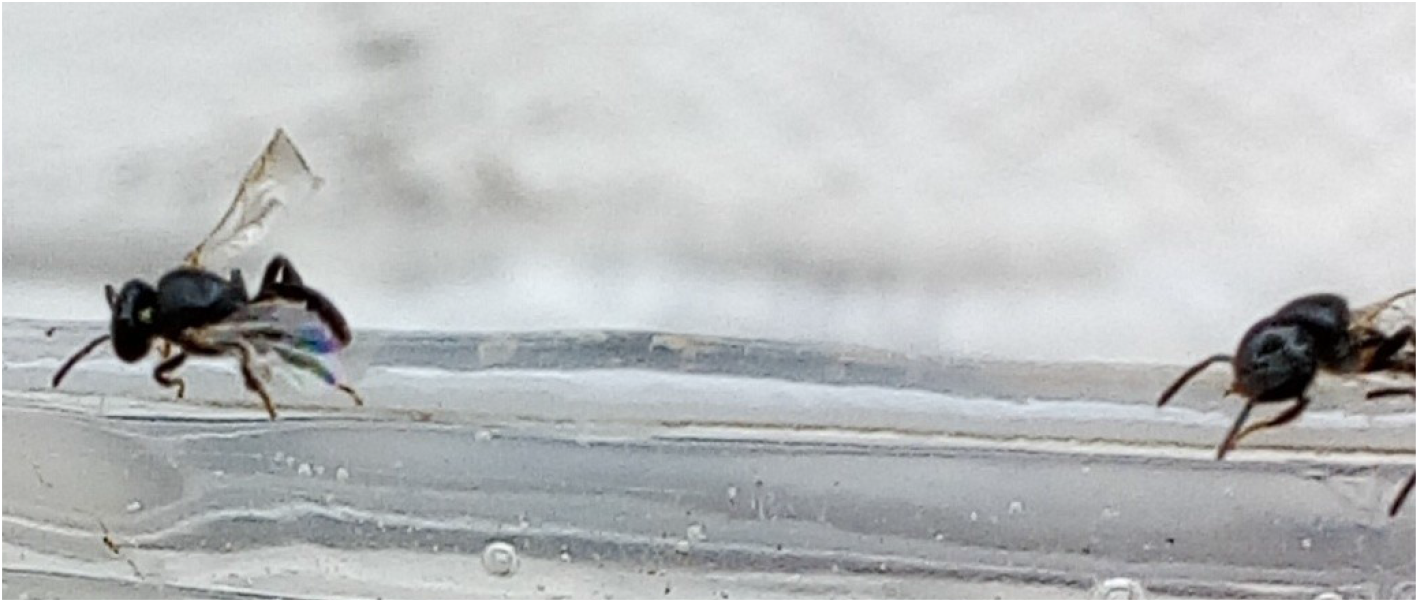
*H. ruspolii* forager exhibiting scent marking behaviour (wing fanning) on an unscented feeder provisioned with nectar.

### Electrophysiological (GC-EAD) responses to natural extracts of forager bees

To identify compounds from both nasonov and tarsal gland extracts of bee foragers to which the chemo-receptors of their antennae are sensitive to, coupled gas chromatography-electro-antennogram detection (GC-EAD) analyses were conducted. This was to establish if meliponine foragers can detect and positively respond to volatile compounds responsible for scent marking behaviour dominant in both nasonov and tarsal gland extracts. Excised antennae (Olsson *et al*., 2013) of foragers from the four meliponine bee species; *Hypotrigona ruspolii, Hypotrigona gribodoi, Meliponula ferruginea* (black) and *Plebeina hildebrandti* were mounted between two capillary glass electrodes filled with saline solution. The electrodes were connected to a high-impedance DC amplifier (Syntech), and the flame ionization detection (FID) and electro-antennographic (EAD) signals were simultaneously recorded on a PC using the program GC-EAD 2000 (Syntech). For each run, 3μl gland extract was injected in split less mode at 50°C onto the column. The Flame Ionization Detector (FID) was heated to 300 °C to detect all compounds. We used an HP-5 column (30 x 0.25 mm ID X 0.25 μm, Agilent, US) with nitrogen (2 ml/min) as the carrier gas. The oven temperature was 50 °C for 2 min and then increased at 10 °C/min to 230 °C. The electro-antennogram (EAG) system was connected to the GC system with a custom, 40 cm heated (250 °C) transfer line. Separate recordings of both EAD and FID signals were done. We replicated EADs with three individual forager antennae from each of the four species. A peak was classified as electro-physiologically active when it coincided with an EAD baseline deflection.

### Extraction of headspace volatiles (nasonov and tarsal glands) for chemical analyses

Headspace volatiles from both nasonov and tarsal glands from ten foraging bees were routinely extracted using the protocol described by Jarau *et al*., (2006). Glands were dissected by excising the 6^th^ and 7^th^ abdominal tergite region (nasonov gland) between the tarsus and metatarsus region (tarsal gland) in sterile saline solution and soaking in lml of pentane for 24 hours at room temperature (24°C), thereafter evaporating the solvent under a gentle stream of nitrogen gas to adjust 100μl per pair of glands (e.g., 10 nasonov glands in 500 μL pentane /10 tarsal glands in 500 μL pentane), thus 100μl of the pooled extracts corresponded to the gland content of one individual bee (one bee equivalent). Extracts were stored in -20°C until ready to use for chemical analyses. A pure pentane control was subjected to similar evaporation process.

### Chemical Analyses

Coupled gas chromatography/mass spectrometric (GC/MS) analysis was carried out on an Agilent Technologies 7890A gas chromatograph equipped with a capillary column HP-5 MS (30 m × 0.25mm ID ×0.25μm film thickness) and coupled to a 5795C mass spectrometer. An aliquot (1 μl) of the gland extracts from different species was injected in split less mode (Inlet temperature = 250 °C, Pressure = 12.1 psi), and helium was used as the carrier gas at 1.0 ml/min. The injector port was maintained at 280 °C. The oven temperature was then held at 35°C for 5 min, increased to 280 °C at 10 °C/min, and then held at 280 °C for 5 min. Mass spectra were recorded at 70 ev. All the alkanes, alkenes. ethers, alcohols, organic acids, esters and aldehydes were identified by comparing their retention times and mass spectral data with those recorded from the NIST 08 spectral library and by co-injection with authentic standards, while the alkenes and aldehydes were identified by using EI diagnostic ions (El-Sayed, 2009). For compound quantification, peak areas were compared to an external standard corresponding to 5ng/μl of 2-heptanol.

### Chemicals

Authentic chemical standards (>95 % purity by GC) (*E)-β* Farnesene, (*Z)-β* Farnesene, Nerolidyl acetate <E>, Bergamotene<α-trans>, Sesquisabinene, Humulene<alpha>, Myrcene, Limonene, Longipinene, Bisabolene<(Z)-alpha, Sinensal <beta>, Funebrene<beta>, Caryophyllene(*E*), Sesquilavandulol <E>, Butanoate<3-methyl-2-butenyl 2-met>, Sesquiphellandrene<beta->, Ocimene<(Z)-beta->, Clovene<alpha-neo->, Himachalene<alpha->, Farnesol<2Z, 6Z) were purchased from Sigma Aldrich (St. Louis, MO, USA).

### Behavioral experiment 3: Scent marking behaviour on food resources baited with synthetic compound, (*E)-β* Farnesene

Bioassays were conducted with pairs of forager bees (N= 25) originating from four different colonies and species were collected from their respective nest entrances while returning from foraging and then immobilized on ice for approximately five minutes to minimize the possibility of the bees producing any alarm pheromones. Exposure to food sources baited with a synthetic form of the dominant compound identified from the nasonov and the tarsal glands: (*E)-β* Farnesene was carried out respectively. Initiation of scent marking behaviour in response to the synthetic compound were conducted in a dual choice test bio-assay Perspex platform measuring 13 x 5.7cm and sealed with a glass lid (**Fig 3**). An aliquot of this synthetic pheromone (25μl) was dispensed round a food source placed onto a filter paper (Whatman No.1) which was placed on one side of the bio-assay chamber while the other chamber was provisioned with an untreated food resource (positive control).

**Figure 3:**
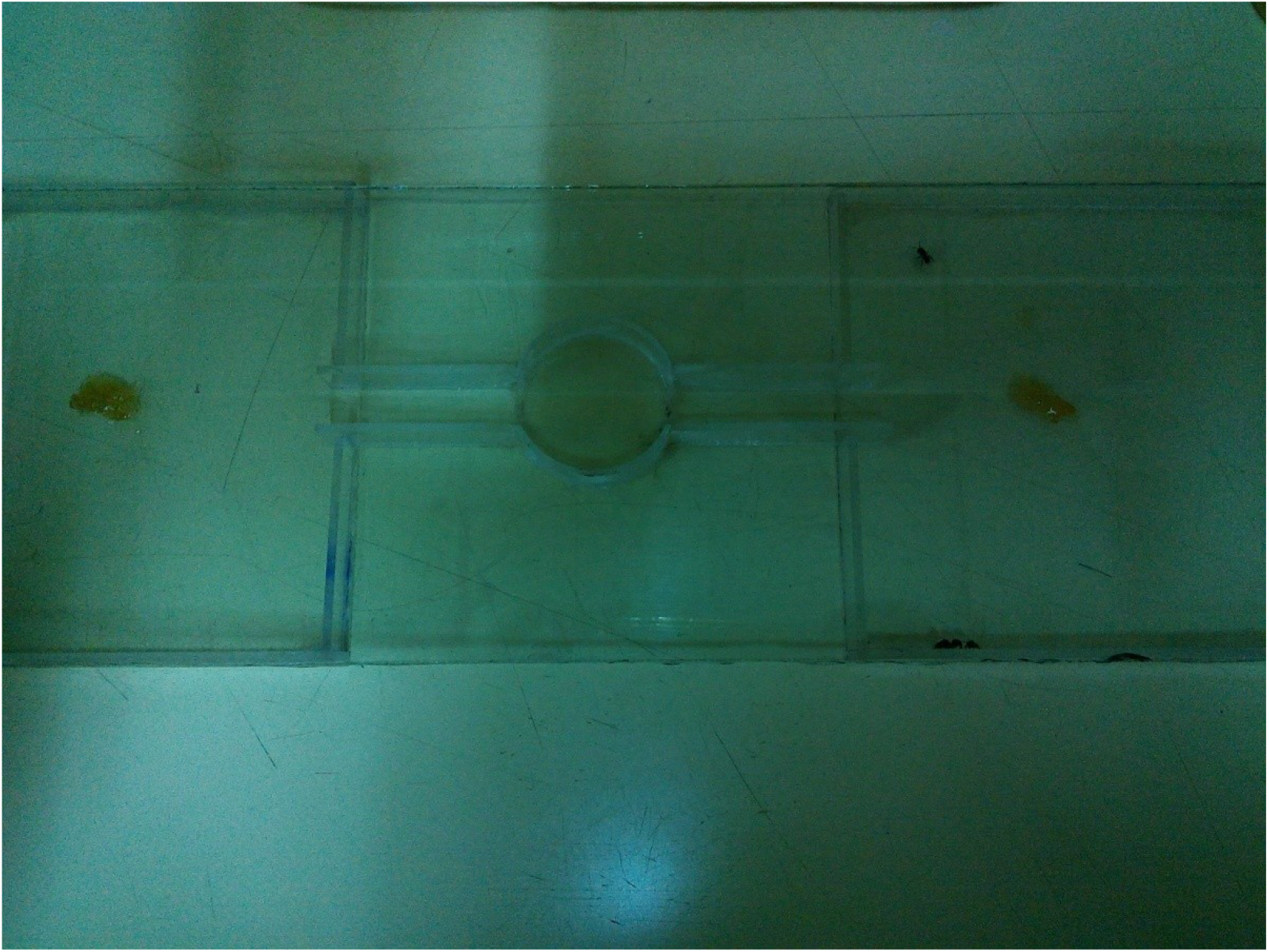
Dual choice test bio-assay Perspex platform provisioned with both baited (treatment) and un-baited food resource (positive control).

**Figure 4a:**
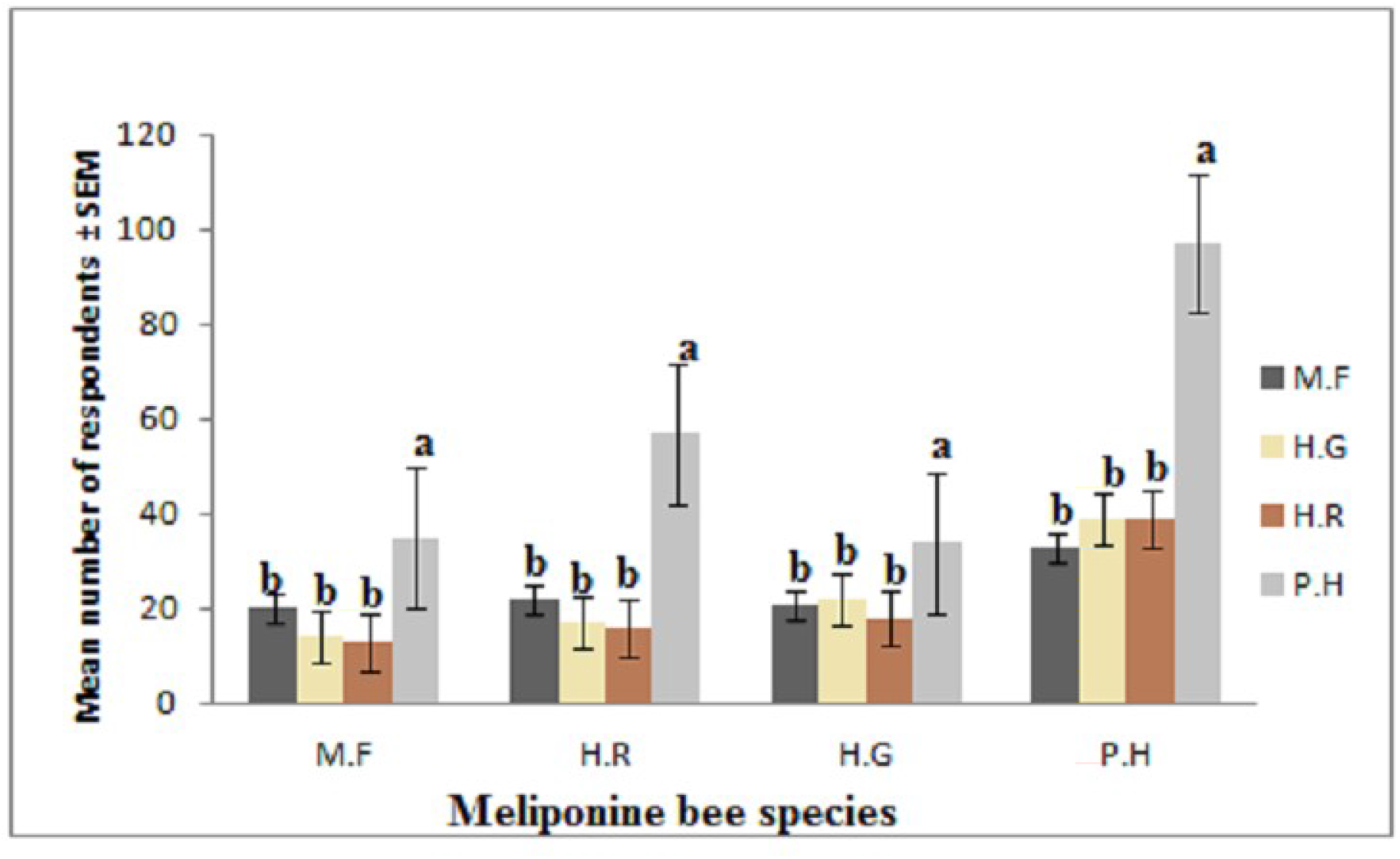
Mean number of individuals recruited to food sites which have been baited with nasonov gland extracts from the respective species. *M.F: *Meliponula ferruginea*, H.R: *Hypotrigona ruspolii*, H.G: *Hypotrigona gribodoi*, P.H: *Plebeina hildebrandti*.

**Figure 4b:**
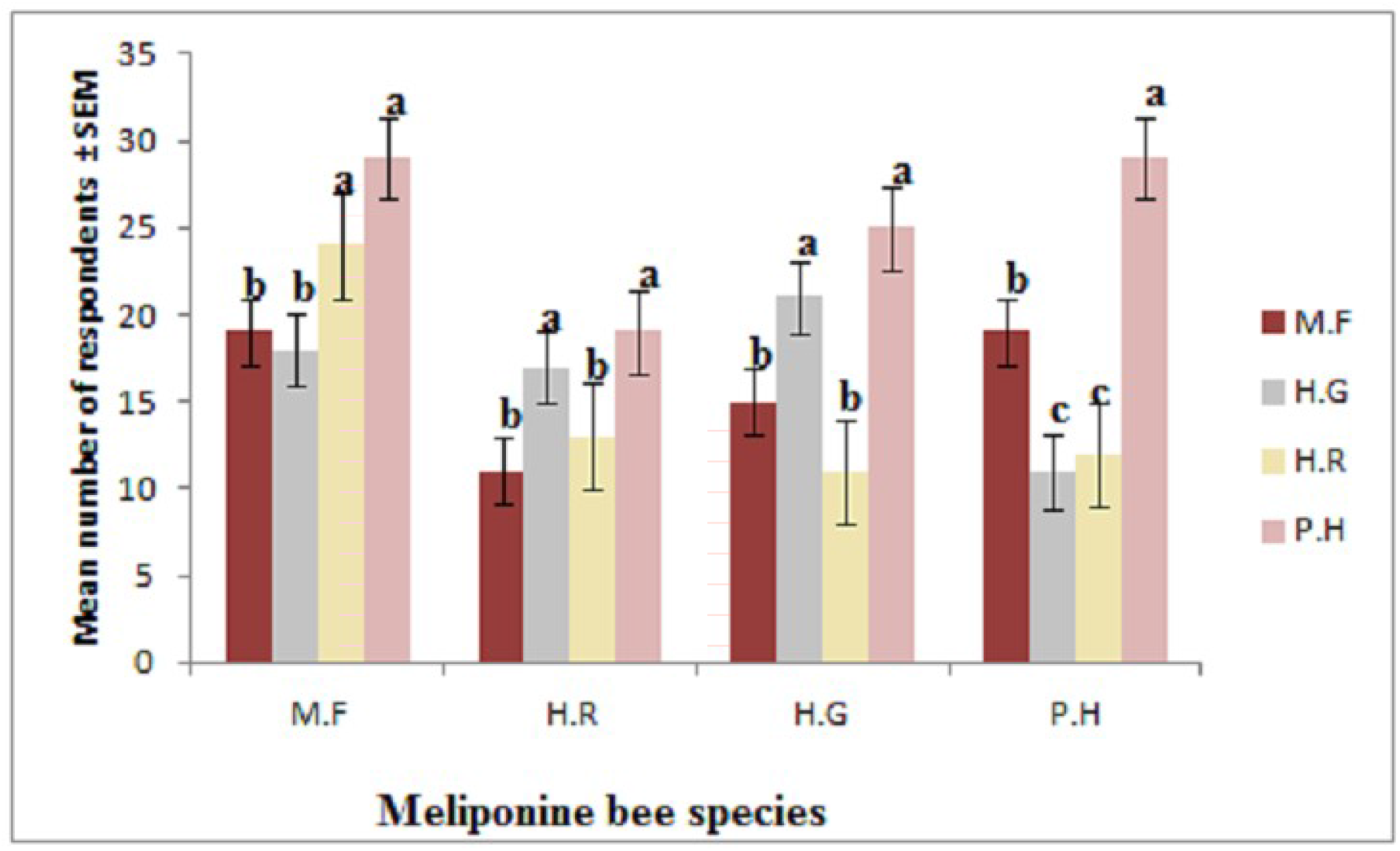
Mean number of individuals recruited to food sites which have been baited with tarsal gland extracts from the respective species*M.F: *Meliponula ferruginea*, H.R: *Hypotrigona ruspolii*, H.G: *Hypotrigona gribodoi*, P.H: *Plebeina hildebrandti*.

### Statistical Analyses

The foraging pattern of individual foragers on each food resource was also analyzed using descriptive statistics. Student Newman Keuls (SNK) tests were used to check for significant effects on foraging behavior to a preference of either treatment (un-baited and baited food sources).

Data from scent marking behavior by all four meliponine bees’ species was subjected to one sample chi-square test by testing for significant differences when exposed to natural extracts of both nasonov and tarsal glands and the tested synthetic compound:

In order to compare the gland composition of trail pheromones of the four different species, the relative peak areas of both nasonov and tarsal gland compound constituents of *Hypotrigona gribodoi, Hypotrigona ruspolii, Meliponula ferruginea* (black) and *Plebeina hildebrandti* were calculated, then further subjected to log-transformed data to Kruskal-Wallis ANOVA test. All statistical analyses were carried out using Sigmaplot V 11.0 statistical software (Systat Software, San Jose, CA 2011).

## RESULTS

### Behavioral experiments 1, 2 and 3: Foraging and scent marking behaviour on food resources

Significant differences were observed in the foraging patterns of each of the four bee species on collected resources (nectar, pollen and water) between 11:00 hours and 14:00 hours; *Meliponula ferruginea* (black) (F_3,116_ =5.61, P<0.001), *Hypotrigona gribodoi* (F_3,116_ =6.46, P<0.001), *Hypotrigona ruspolii* (F_3,116_ =2.81, P=0.042) and *Plebeina hildebrandti* (F_3,116_ =4.19, P=0.007). In all four species, the total number of bees landing and initiating scent marking progressed with increasing foraging hours. Foraging activity peaked between 11:00 hours and 14:00 hours as 70% of all foraging bouts gradually declined after this observation period. *Meliponula ferruginea* (black) species showed the highest foraging activity on both baited and un-baited nectar sources, as workers began landing on the feeders as from 11:05hours and peaked at 13:00hours, while *Hypotrigona gribodoi, Hypotrigona ruspolii* and *Plebeina hildebrandti* all foraged till much later, signifying similar commencement of foraging but having peak periods which lasted until 15:00hours. In general, the collection of nectar started to decrease after this peak period until cessation. Nectar was always the most collected resource *Meliponula ferruginea* (black) (N=220), *Hypotrigona gribodoi* (N=117), *Hypotrigona ruspolii* (N=124), *Plebeina hildebrandti* (N=109, throughout the whole observational period, while water was the second most collected resource: *Meliponula ferruginea* (black) (N=101), *Hypotrigona gribodoi* (N=97), *Hypotrigona ruspolii* (N=94), *Plebeina hildebrandti* (N=71, followed lastly by pollen: *Meliponula ferruginea* (black) (N=84), *Hypotrigona gribodoi* (N=60), *Hypotrigona ruspolii* (N=61), *Plebeina hildebrandti* (N=73).

The foraging activity for pollen followed the same sequence across all four species, but with no significant difference in activity. Two species showed similar foraging peaks for this resource from 12:00 hours for 50% of the observational period; *Hypotrigona gribodoi* and *Hypotrigona ruspolii* foragers, which was characterized by constant number of bees landing on the feeders with pollen and eventually decreased as the day progressed compared to *Meliponula ferruginea* (black) and *Plebeina hildebrandti*.

Notable however was the foraging pattern for water which was observed to be more regular after 13:00hours.

### Chemical and electrophysiological analyses

Chemical analyses of both nasonov and tarsal gland extracts demonstrated that the trail pheromone of *Plebeina hildebrandti, Hypotrigona gribodoi, Hypotrigona ruspolii* and *Meliponula ferruginea* could be potentially produced by nasonov glands but mechanically deposited on any surface through the tendon retractor claws located on the hind legs, based on scent marking observations. Four dominant compounds were identified from the nasonov gland extracts (**Figure 5a**) and two dominant compounds from the tarsal gland extract (**Figure 5b**) which were all sesquiterpenes.

**Figure 5a:**
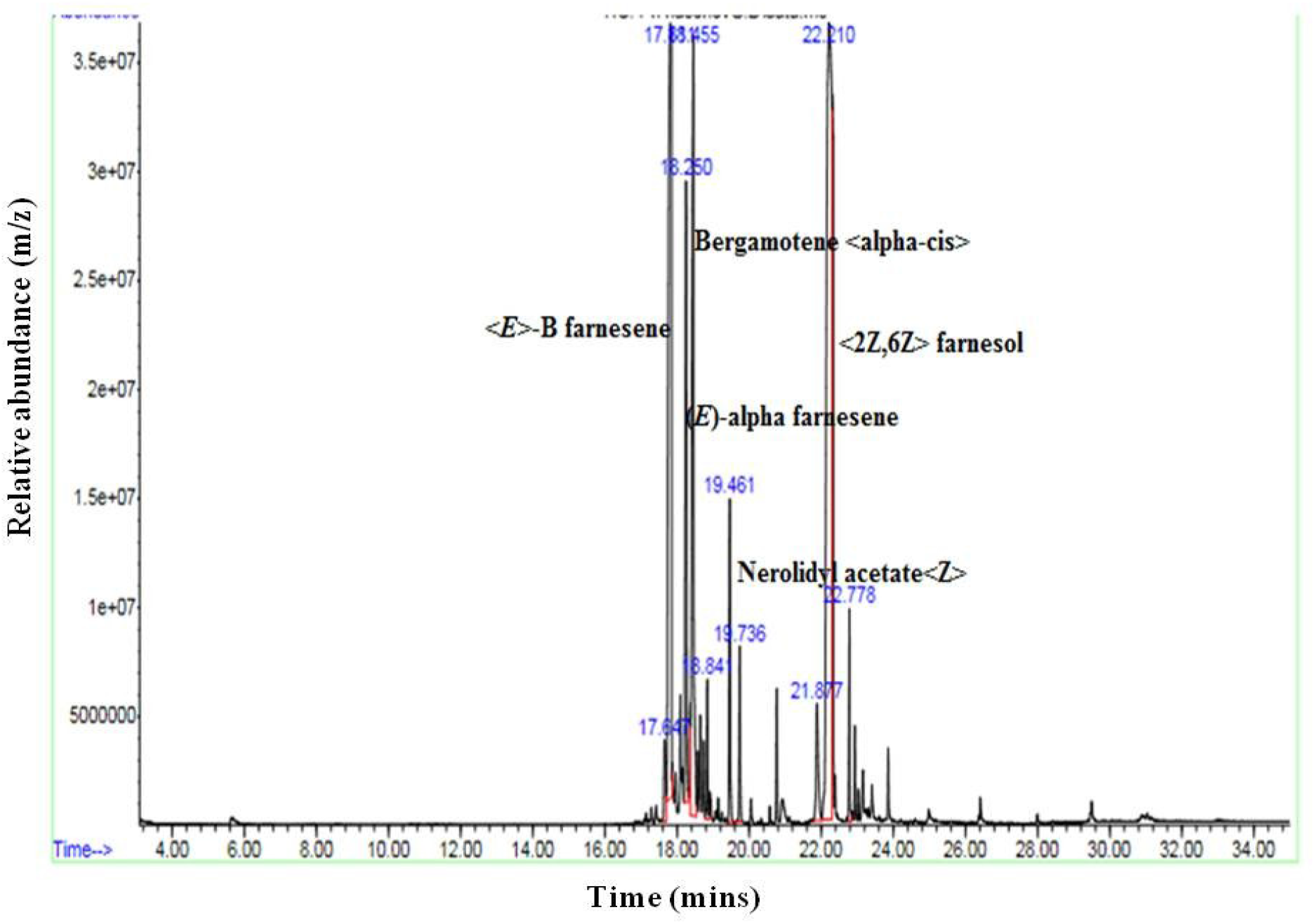
Mass spectrum showing dominant compounds identified from the nasonov epithelial gland extract of a meliponine bee species, *Hypotrigona ruspolii*.

**Figure 5b:**
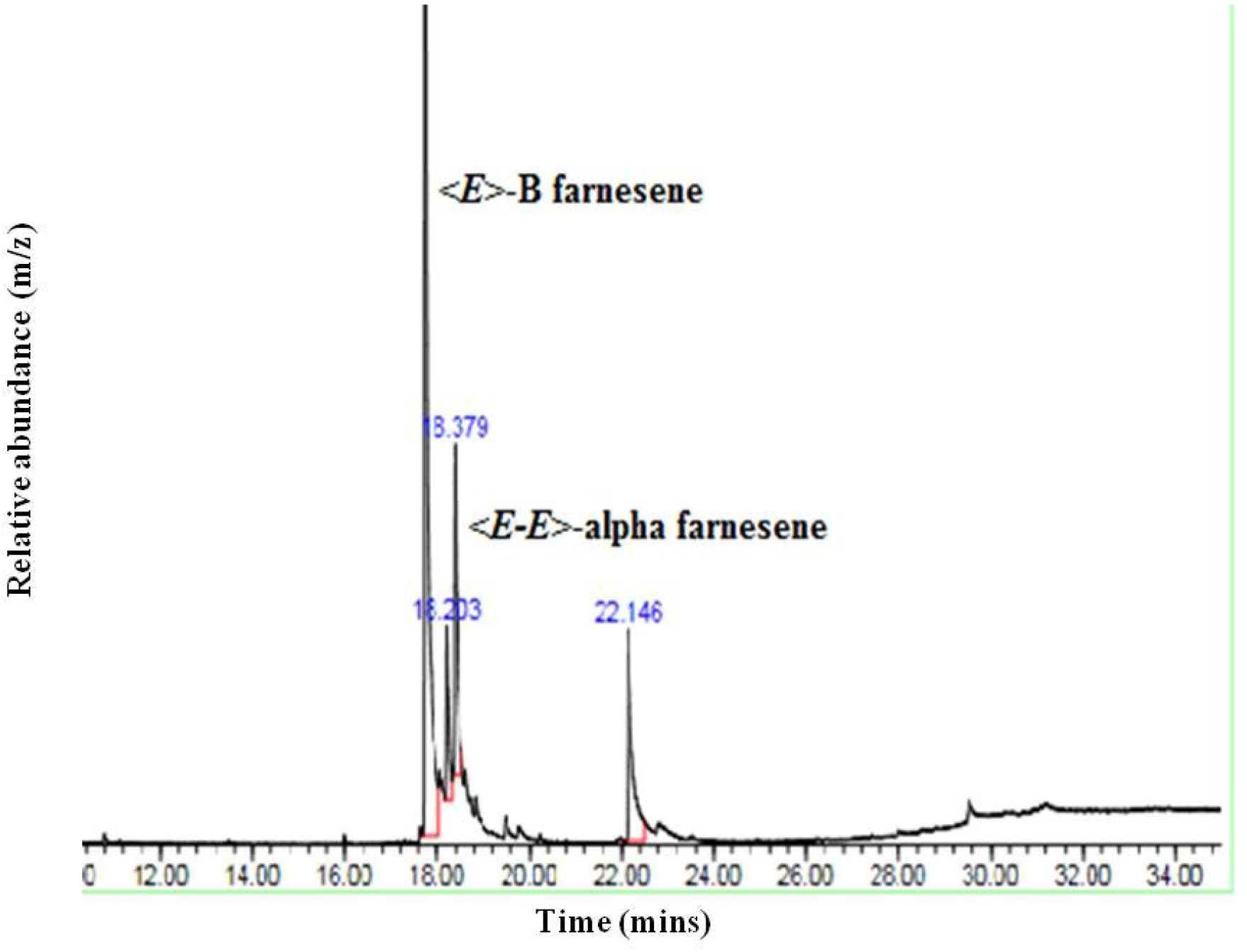
Mass spectrum showing dominant compounds identified from the tarsal gland extract of a meliponine bee species, *Hypotrigona ruspolii*.

GC-EAD analyses done with 30 worker bee antennae revealed one peak that elicited consistent responses of the chemoreceptor’s in more than 30% of the trials. These peaks correspond to the compound: (*E*)-β Farnesene. The physiological activity of (*E*)-β Farnesene was verified in subsequent GC-EAD runs with its synthetic derivative.

### Bio-assays with synthetic compounds

#### Scent trail bioassays with synthetics

To test whether the physiologically active compounds from both nasonov and tarsal glands constitute the behaviorally active trail pheromone of these species, a further set of trail bioassays was conducted. Experimental trails were baited with (*E*)-β Farnesene which was the compound with the most dominant peak in both the nasonov and tarsal gland secretions of foragers collected from the four species respectively; *Hypotrigona gribodoi, Hypotrigona ruspolii, Meliponula ferruginea* (black) and *Plebeina hildebrandti* respectively.

A significantly higher proportion of foragers from the four species were attracted and recruited additional foragers to food resources baited with natural extracts from their own nasonov glands: *M. ferruginea* (black) (t = 4.097 df =58, P< 0.001), *Hypotrigona ruspolii* (t = 0.633, df = 58, P = 0.005), *Hypotrigona gribodoi* (t = 2.64, df = 58, P= 0.004) and *Plebeina hildbrandti* (t =12.92, df = 58, P<0.001) over the control (un-baited food resource), (F = 95.77, df =4, 145, P < 0.001). This similarly occurred when compared to food sources baited with natural extracts from their own tarsal glands or from other species; with no significance preference for any species: *M. ferruginea* (black) (t = 2.41, df = 58, P= 0.011), *Hypotrigona ruspolii* (t = 2.49, df = 58, P = 0.015), *Hypotrigona gribodoi* (t = 2.52, df = 58, P = 0.014) and *Plebeina hildebrandti* (t = 2.85, df = 58, P = 0.006) over the control (un-baited food resource) (F = 1.22, df = 4, 145, P=0.304), as no significant differences was observed between respective treatments. The synthetic compound, (*E*) - β Farnesene was significantly as attractive to foragers of the four species when compared to the natural nasonov gland extract but not natural tarsal gland extracts ((*E*) – β Farnesene: (F = 19.01 df = 4, 145, P<0.001), nasonov gland extract: (F = 95.77, df =4, 145, P < 0.001), tarsal gland extract: (F = 1.13, df = 4, 145, P=0.304).

## Discussion

The results of our bio-assays show that some African meliponine bee species carry out scent marking at food sources and trail pheromones of these four species may exclusively be produced in the foragers’ exocrine glands. This is in accordance with recent studies conducted with *Scaptotrigona pectoralis, Geotrigona mombuca, Trigona recursa* and *Trigona spinipes* (Jarau *et* al., 2000, 2003b; 2006; 2010; Stangler *et* al., 2009; Reichle *et* al., 2013) and further disclaims the long assumed role of mandibular gland secretions for scent trail marking in meliponine bees species (Lindauer and Kerr, 1958, 1960; Kerr *et* al., 1963; Nieh *et* al., 2003; 2004; Kuhn-Neto *et* al., 2009; Lichtenberg *et* al., 2011).

The compound from nasonov gland extracts detected by the chemo-receptors on the foragers’ antennae from these four species belongs to the chemical class of terpenoids. Gas chromatographic analysis had shown that this compound, (*E)*-β Farnesene constitutes a dominant part of the trail pheromone in these species. However, the natural nasonov gland extract was more attractive to recruited foragers, compared to the singular sesquiterpene compound, (*E)*-β Farnesene. The most obvious reason for this may be that this physiologically active compound may be in-complete as a synthetic pheromone trail bouquet, which has been shown to contain varied amounts of geraniol and citral in some studies (Jara*u et* al., 2003b; Stangle*r et* al., 2009; Hrnci*r et* al., 2016).

This study therefore adds to the existing list of known trail pheromone compounds used by meliponine bee species, and it can be assumed that the terpenyl esters identified from nasonov or tarsal gland extracts of other trail laying species may constitute their respective unique trail pheromones. Indeed, the chemical similarities between these compounds such as the terpenyl esters in these meliponine bees are also used as marking compounds by some solitary bees and bumblebees by depositing carboxylic acid alkyl esters on twigs or leaves for mating purposes (Bergstrom, 2008).

In this present study, a generality of compounds from the terpenyl esters group in the trail pheromones of *Plebeina hildebrandti, Meliponula ferruginea* (black), *Hypotrigona gribodoi* and *Hypotrigona ruspolii* in terms of composition, was sufficient in triggering trail-following behavior. This conclusive finding, reveals that foragers are significantly attracted to food sources baited with nasonov gland extracts prepared from their nest-mates over foragers of a foreign colony, and this may be further explained by the differences in the relative proportions of trail pheromone components of foragers from these different species.

Though there seemed to be some minute disparity in the scent marking components of these bee species, which could either be linked to their morphology such as body and gland size which could influence relative abundance of these scent marking compounds. It was observed that the gland components of larger sized bees, *Plebeina hildebrandti* was dominated by larger amounts of terpenoids compared to much smaller sized species, *Hypotrigona gribodoi, Hypotrigona ruspolii* and *Meliponula ferruginea* (black). It may be that these smaller sized bees significantly make use of other compounds such as cuticular hydrocarbons to lay trails, and this has been observed from certain studies suggesting that cuticular hydrocarbons could also provide and function as footprint cues in social wasps and some bee species to recognize their nest entrance at close range (Soroke*r et* al.,1998). Similarly, these same footprint hydrocarbons are informative to foraging bees, and are readily used to discriminate against either visited or already depleted food sites (Goulso*n et* al., 2000, 2002; Bart*h et* al., 2008; Jara*u et* al., 2012). Although this discriminating behavior originally was believed to be based on active deposition of lipid “scent-marks” by bees, two recent studies suggested that these chemicals are deposited wherever the bees walk, and were used as footprint cues rather than pheromonal signals. This sheds more light to the dual functionality that cuticular hydrocarbons may play in communication mechanisms. *Bombus terrestris* workers were reported to deposit a similar range of compounds, mostly long chain alkanes and alkenes, in essentially similar concentrations at food, nest, and neutral sites (Goulso*n et* al., 2000). These findings suggest that these hydrocarbon marks are deposited involuntarily, regardless of the present behavioral context of the foraging bee (Nieh and Roubik, 1995; Schmid*t et* al., 2003; Hrnc*ir et* al., 2004). Recently, Holldobler *et al*. (2004) reported that the preference of *P. rugosus* foragers for food-sites marked by their nest mates over food-sites marked by foreign con-specific workers is likely due to similar gland secretions, which contain nest-specific patterns of volatiles (mainly hydrocarbons and esters) deposited in addition to the abdominal gland content. A colony-specific effect of abdominal extracts in releasing trail following behavior was demonstrated in *Lasius neoniger* (Traniello, 1980) but by contrast, the actual trail pheromone extracted from workers abdominal region was not specific in initiating scent marking behavior in other closely related species, *Lasius japonicus* and *Lasius nipponensis*. However, colony specificity was added to these trails by footprint hydrocarbons deposited by these workers along their own trails (Akino and Yamaoka 2005; Akino *et al*., 2005).

## Conclusion

Regardless of the dominant presence of the sesquiterpene *E*-β farnesene in *Plebeina hildebrandti, Meliponula ferruginea* (black), *Hypotrigona gribodoi* and *Hypotrigona ruspolii* glandular extracts and its use as a scent marking cue, it is likely that foragers are able to detect and distinguish scent trails deposited by workers of both same and foreign species by recognizing other compounds secreted in minute quantities which could function as a pheromone bouquet. Avoiding foreign scent trails appears advantageous to these bees because they reliably indicate the location of an already visited food source which could help in avoiding both competition and conflicts at food sources between foragers of different species. This is of particular importance for the survival of less aggressive meliponine bee species. A forager’s ability to discriminate between trails laid by a different forager of another species is most likely based on the recognition of additional but minute compounds in their specific pheromone bouquets.

When foragers of different species meet at food sites, they become entangled in fierce fights, which usually lead to the death of many individuals (Johnson and Hubbell, 1974; Hncrir *et al*., 2004). Hence, by limiting aggressive encounters between foragers of different species, the loss of large numbers of workers could be avoided and the colonies’ fitness maintained. This could potentially be a mechanism for resource partitioning and competition avoidance between con-specific and hetero-specific foragers, which occurs when foragers from different colonies have been domesticated and scout for food sources in overlapping foraging areas such as green houses.

